# Intergenerational effects of dietary restriction on insulin/IGF signaling and reproductive development

**DOI:** 10.1101/342956

**Authors:** James M. Jordan, Jonathan D. Hibshman, Rebecca E. W. Kaplan, Amy K. Webster, Abigail Leinroth, Ryan Guzman, Colin S. Maxwell, Elizabeth Anne Bowman, E. Jane Albert Hubbard, L. Ryan Baugh

## Abstract

The roundworm *C. elegans* transiently arrests larval development to survive extended starvation (*1*), but such early-life starvation reduces reproductive success (*2, 3*). Maternal dietary restriction (DR) buffers progeny from starvation, increasing reproductive success (*4*). It is unknown why early-life starvation decreases reproductive success and how maternal diet modifies this process. We show here that extended starvation in first-stage (L1) larvae followed by unrestricted feeding results in a variety of abnormalities in the reproductive system, including *glp-1/*Notch-sensitive germ-cell tumors and uterine masses that express neuronal and epidermal markers. We found that maternal DR reduces the penetrance of starvation-induced abnormalities, including tumors. Furthermore, we show that maternal DR reduces insulin/IGF signaling (IIS) in progeny, and that *daf-16*/FoxO and *skn-1*/Nrf, transcriptional effectors of IIS, are required in progeny for maternal DR to suppress abnormalities. *daf-16*/FoxO activity in somatic tissues is sufficient to suppress starvation-induced abnormalities, suggesting cell-nonautonomous regulation of reproductive system development. This work reveals complex inter- and intra-generational effects of nutrient availability mediated by IIS with consequences on developmental integrity and reproductive success.

**One Sentence Summary:** Intergenerational effects of diet on IIS

## Main Text

To determine how extended larval starvation compromises reproductive success, we compared early adult worms starved for 8 days as L1-stage larvae to control adults that were starved briefly, hereafter referred to as “starved” and “control”, respectively. Approximately half of the starved worms displayed prominent abnormalities in their reproductive system despite being well fed after the starvation period (Fig. 1A-C). These abnormalities were highly variable, both among worms and between gonads in an individual worm. The most common abnormalities fell on a spectrum ranging from gonads enlarged with proliferative germ cell nuclei (detected as P*mex-5*::H2B::mCherry-positive, “*mex-5*^+^”) to uterine masses consisting of largely P*mex-5*::H2B::mCherry-negative cells (“*mex-5*^-^”, Fig. 1C). The *mex-5*^+^ gonad enlargements resembled proximal germ cell tumors similar to those caused by “latent niche” signaling (*5, 6*), and the *mex-5*- uterine masses were disorganized and appeared to contain differentiated cells. Indeed, the majority of uterine masses expressed *elt-1* and *unc-119* reporter genes, embryonic markers of epidermis and neurons, confirming somatic differentiation (Fig. 1D and E). We observed similar abnormalities in three wild isolates subjected to 8 d L1 starvation (Fig. S1), suggesting that these abnormalities were not an artifact of the laboratory strain. Individuals with abnormalities produced smaller broods (Fig. 1F), consistent with developmental abnormalities limiting reproductive success.

**Fig. 1.**
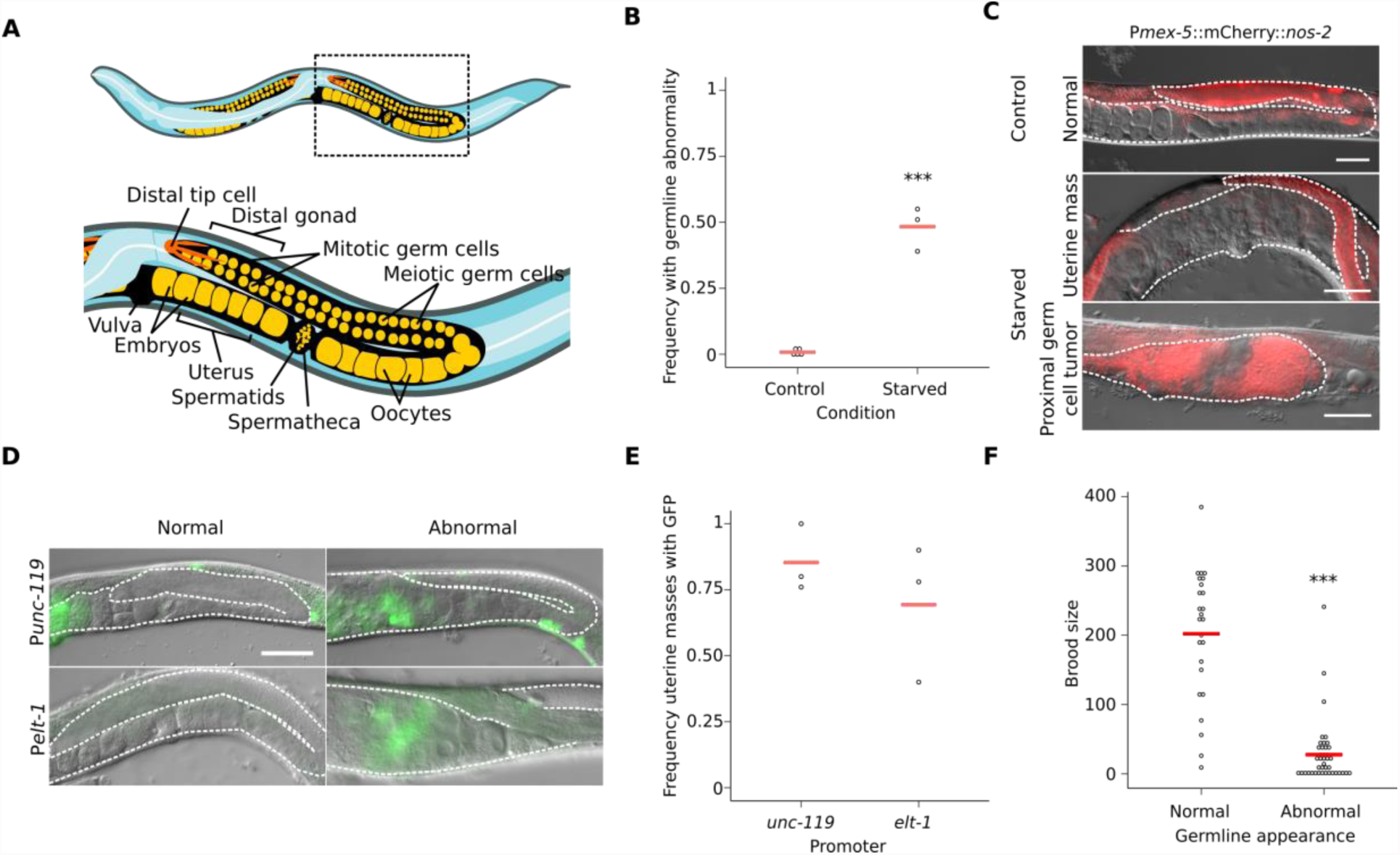
Early-life starvation followed by *ad libitum* feeding results in germline abnormalities. . (A) Cartoon depicting organization of posterior gonad arm of an adult *C. elegans* hermaphrodite. Boxed area is enlarged to show region assessed for germline abnormalities. The cartoon is not to scale and does not depict all germ cells. (B) Circles indicate biological replicates scoring at least 50 worms per condition per replicate. (C) Representative images of adult gonad arms from given conditions. (D) Representative images of adult gonad arms after L1 starvation with (“Abnormal”) or without (“Normal”) uterine masses. (C and D) Gonad is outlined with a white dashed line. Animals are oriented as in A. Scale bars are 50 microns. (E) Circles indicate the frequency of uterine masses that are GFP-positive. At least 20 masses were scored per condition per replicate. (F) Circles indicate individual brood sizes from two biological replicates of 31 and 35 starved worms total. (B and F) ***p < 0.001; t-test. (B and E) Cross bars reflect the mean.

The Notch receptor GLP-1 and the RNA-binding protein GLD-1 regulate germline differentiation (*7*). *glp-1* activity maintains germline stem cells by preventing differentiation, and gain-of-function mutants develop germ cell tumors (*8, 9*). We found that the germline abnormalities we see in starved wild-type worms resemble proximal tumors seen in *glp-1(ar202)* gain-of-function mutants, and starvation increased penetrance of tumor formation in the mutant (Fig. 2A and B)(*8*). Interestingly, the presence of abnormalities was sensitive to *glp-1* dosage, since few animals heterozygous for the *glp-1(e2072)* loss-of-function mutation (*10*) displayed germline abnormalities following L1 starvation (Fig. 2B). *gld-1* promotes entry into meiosis and differentiation, and class A loss-of-function mutants develop germ cell tumors (*11*). In our control conditions, early-adult *gld-1(q485)* null mutants developed proximal germ cell tumors but heterozygous mutants did not, as expected. In worms subjected to 8 d L1 starvation, however, heterozygous mutants developed tumors and the size of the tumors in homozygous mutants was greatly enlarged (Fig. 2C and D). *gld-1* mutants do not perform physiological apoptosis (*12, 13*). Thus, the absence of large tumors in control *gld-1* mutants and our *glp-1* results suggest that extended L1 starvation causes hyperproliferation of germ cells, disrupting coordination of germ line development. Based on our observation of proliferative germ cells in the proximal gonad (adjacent to the vulva), we believe that extended starvation causes heterochronically discordant somatic and germline development resulting in signaling from the “latent niche” (modeled in Fig. 2E)(*5, 6*).

**Fig. 2.**
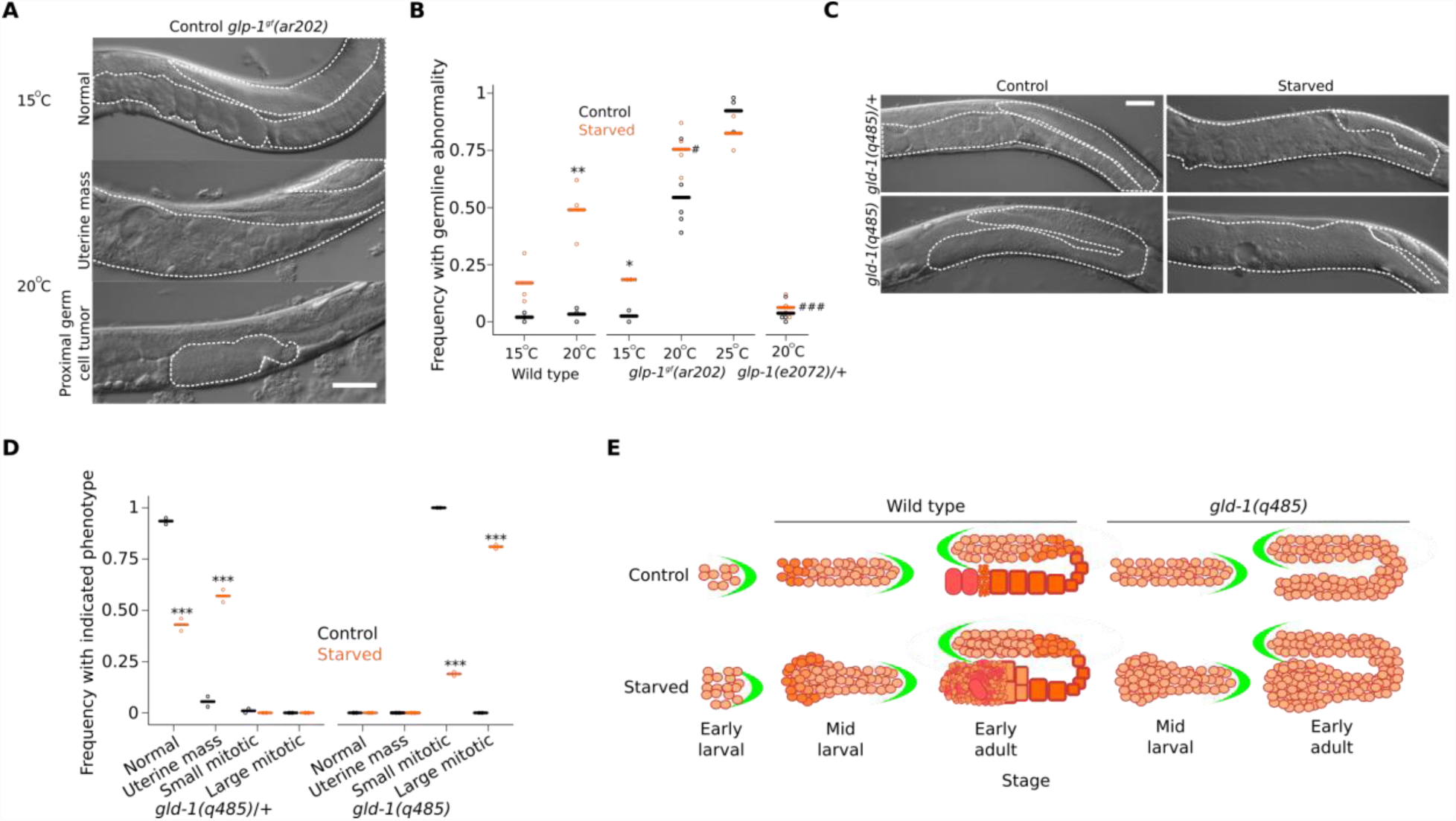
Early-life starvation followed by *ad libitum* feeding results in the formation of *glp-1*/Notch-sensitive germ-cell masses. (A) Representative image of *glp-1^gf^(ar202)* from control conditions at permissive and semi-permissive temperature. (B) Circles indicate biological replicates scoring at least 40 worms per condition per replicate. *p < 0.05, ** < 0.01; t-test between control and starved. #p < 0.05, ### < 0.001; t-test between starved wild type. (C) Representative images of adult *gld-1(q485)* mutants from given conditions. (D) Gonad arms were assigned to four classes as indicated and biological replicates scoring at least 30 worms per condition each are plotted as circles. ***p < 0.001; t-test between frequencies of control and starved abnormalities of given type. (B and D) Cross bars reflect the mean. (E) Model depicting how early-life starvation impairs reproductive development. Green crescent is the distal tip cell (DTC). Undifferentiated cells (light orange circles) enter meiosis and begin to differentiate (darker orange circles) at a specific distance from the DTC before completing differentiation as sperm (small darkest orange circles) and oocytes (rectangles). Fertilized embryos are depicted as red ovoids. Differentiation of oocytes is defective in *gld-1* mutants, therefore hyperproliferation is evident in adulthood of starved animals (Fig. S3D). Early aberrant development following L1 starvation likely alters spatiotemporal signaling between the somatic gonad and germ cells resulting in variable abnormalities in wild-type animals.

Maternal DR buffers progeny reproductive success from the costly effects of L1 starvation (*4*), so we wondered whether maternal DR reduces the incidence of germline abnormalities caused by extended L1 starvation. We acquired progeny from parents fed *ad libitum* (AL) or DR using well-controlled liquid-culture conditions (*4, 14*) and starved them for 8 d as L1 larvae. Fewer DR progeny developed germline abnormalities after starvation than AL progeny (Fig. 3A), revealing an intergenerational effect of DR on pathological consequences of early-life starvation.

Analysis of transcript abundance by RNAseq revealed intergenerational effects of maternal DR on progeny gene expression. 114 genes were differentially expressed on the first day of L1 starvation in DR progeny compared to AL progeny (FDR < 0.1; Table S1). Gene Ontology (GO) term enrichments included terms related to IIS, including “innate immune response” and “regulation of dauer entry” (Table S2). In particular, expression of the only known IIS receptor *daf-2* was decreased in DR progeny (Fig. 3B). DAF-2/InsR antagonizes function of the FoxO transcription factor DAF-16 in response to IIS (*15*), and regulatory targets of *daf-16*/FoxO define a transcriptional signature of reduced IIS (*16*). Despite the relatively small number of differentially expressed genes, genes up-regulated in DR progeny were significantly enriched for Class I *daf-16* targets (activated by *daf-16*; p < 0.05), and down-regulated genes were significantly enriched for Class II targets (repressed by *daf-16*; p < 0.001) (Table S3), consistent with reduced IIS in DR progeny. DAF-2/InsR regulates localization of DAF-16, with DAF-16 shifting from cytoplasmic to nuclear when IIS is reduced (*15*). GFP::DAF-16 displays strong nuclear localization during L1 starvation (*17*), but appeared even more completely localized to nuclei in DR progeny (Fig. 3C). GFP::DAF-16 was relatively more cytoplasmic in fed than starved larvae, as expected, but it remained relatively more nuclear in DR progeny (Fig. 3C). These results collectively support the conclusion that maternal DR decreases IIS in progeny.

**Fig. 3.**
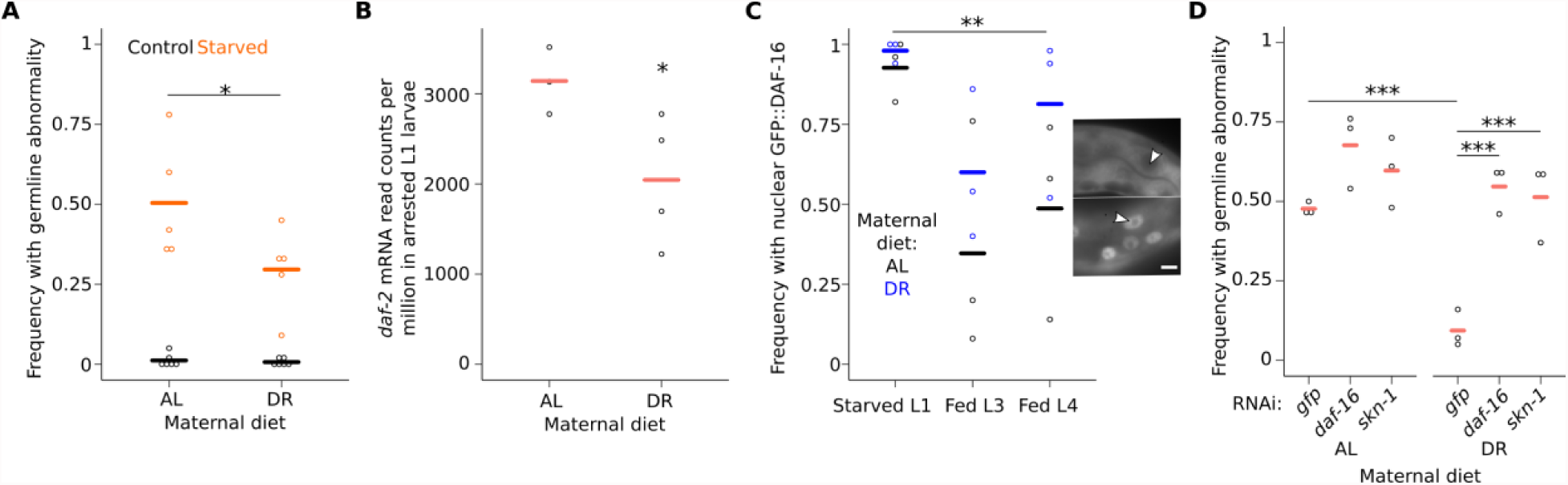
Maternal dietary restriction reduces progeny insulin-like signaling, mitigating early-life starvation-induced germline abnormalities. (A) Circles indicate biological replicates scoring an average of 49 progeny from *ad libitum-*fed (AL) or dietary restricted (DR) parents per replicate. *interaction p-value < 0.05; two-way analysis of variance (ANOVA). (B) Circles indicate *daf-2* mRNA read counts per million (CPM) in arrested L1 progeny of AL or DR parents. *FDR < 0.1. (C) Circles indicate biological replicates scoring 50 worms per condition per replicate. **p < 0.01; paired t-test of AL vs. DR at all timepoints. Insets are representative images of L1 larvae with cytoplasmic GFP::DAF-16 localization (top) or nuclear GFP::DAF-16 localization (bottom) with white arrows indicating intestinal nuclei. Scale bar is 5 microns. (D) Frequency of starved progeny from AL and DR parents that developed a germline abnormality when recovered to adulthood on given RNAi (n = at least 50 worms per replicate). ***p < 0.001; t-test. (A-D) Cross bars reflect the mean.

To determine whether intergenerational reduction of IIS caused by maternal DR has functional consequences, we examined the effects of reduced *daf-16*/FoxO and *skn-1*/Nrf, both effectors of IIS (*15*). Disruption of either gene by RNAi did not alter the incidence of abnormalities in AL progeny starved for 8 d as L1 larvae, but they were each required for maternal DR to reduce the frequency of abnormalities (Fig. 3D). These results further support the conclusion that maternal DR reduces progeny IIS, protecting progeny from pathological consequences of extended starvation by promoting the activity of *daf-16*/FoxO and *skn-1*/Nrf.

Fewer starved larvae that recover in low IIS conditions display germline abnormalities. Dramatically fewer starved larvae recovering in DR or in conditions that induce dauer diapause (*18*) (*19*) (followed by recovery from dauer in AL conditions) displayed abnormalities (Fig. 4A and Fig. S2A). These results are consistent with a protective effect of reduced IIS during development after starvation, but other pathways affected by these conditions could also be involved. Mutation of *daf-2*/InsR also suppressed abnormalities (Fig. 4B). Likewise, *skn-1*/Nrf gain-of-function mutants suppressed abnormalities in *daf-16*-independent fashion (Fig. 4C). Though consistent with reduced IIS suppressing starvation-induced germline abnormalities, it is unclear when during the lifecycle these mutations exert their effect. We therefore used RNAi to perturb gene function during development after starvation. RNAi of *daf-2* (or double RNAi of *daf-2* and *gfp*) suppressed the frequency of abnormalities (Fig. 4D and E), demonstrating that reducing IIS during recovery of previously starved larvae suppresses abnormalities. Suppression depended on *daf-16*/FoxO and *skn-1*/Nrf (Fig. 4E), as seen with maternal DR (Fig. 3D). To determine whether these results were due to RNAi depletion of IIS genes in the germline or soma, we performed RNAi in mutants that largely restrict RNAi to these tissues (*20, 21*). The effects of *daf-2* and *daf-16* RNAi were abrogated in a mutant background that restricts RNAi primarily to the germ line but were retained in a mutant background that restricts RNAi primarily to the soma (Fig. 4F), indicating that these genes function in the soma to suppress the observed abnormalities in the reproductive system. Tissue-specific transgenic rescue of a *daf-16* null mutant showed that overexpression of *daf-16* without *daf-2* RNAi is sufficient to suppress abnormalities but that *daf-2* RNAi enhances suppression (Fig. 4G). These results corroborated results using double RNAi (Fig. 4E and F), and they revealed that *daf-16* can function cell-nonautonomously in the epidermis, intestine or neurons to regulate post-starvation germline development. These sites of action have been shown for *daf-16* in regulating developmental arrest and organismal aging (*22-25*), but not germline proliferation where *daf-2* acts germ cell-autonomously in late-larval stages to promote germ cell proliferation (*26*) but acts cell-nonautonomously in the somatic gonad during aging to antagonize germline progenitor cell loss (*27*). Thus, the role of IIS in germline development during recovery from L1 starvation involves a different mechanism. Nonetheless, multiple lines of evidence support the conclusion that reduction of somatic IIS during development suppresses starvation-induced germline abnormalities.

**Fig. 4.**
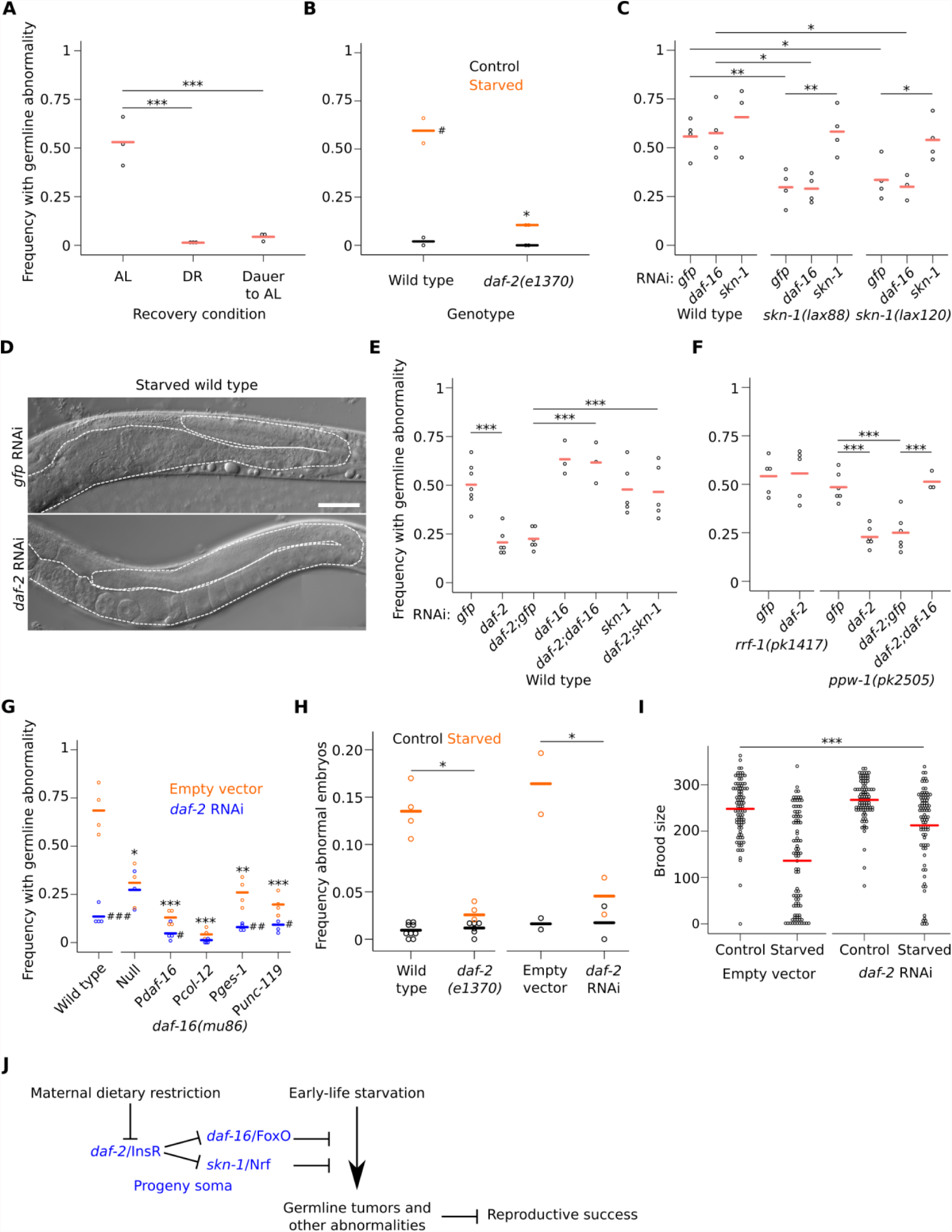
Reducing somatic insulin/IGF signaling during recovery from early-life starvation suppresses germline abnormalities. (A) Wild-type worms were starved for 8 d and recovered in *ad libitum* (AL), dietary restriction (DR) or dauer-forming conditions followed by AL feeding (Dauer to AL). Circles indicate three biological replicates scoring at least 40 worms per condition per replicate. (B) Circles indicate biological replicates scoring at least 38 animals per condition per replicate. (C, E, and F) Circles indicate biological replicates scoring at least 50 starved worms per condition per replicate. (D) Representative image of worms from indicated conditions. Gonad and uterus are outlined with a dashed white line. Scale bar is 50 microns. (G) Tissue-specific rescue of *daf-16.* Circles indicate biological replicates scoring at least 50 worms per condition per replicate. Asterisks indicate comparisons between wild type and the other genotypes. Pound signs indicate comparisons within genotypes between empty vector and *daf-2* RNAi; #p < 0.05, ## < 0.01, ### < 0.001; t-test. *daf-16(mu86)* null mutants do not survive 8 days of L1 starvation, therefore these animals were starved for three days. Other genotypes were starved for 8 d. (H) Circles indicate biological replicates of abnormal embryo frequencies for *daf-2* mutant and RNAi, scoring at least 100 embryos per condition per replicate. (A-C and E-H) *p < 0.05, ** < 0.01, *** < 0.001; t-test. (I) Circles indicate individual brood sizes from five biological replicates of ~18 individuals per condition per replicate. A linear mixed effect model was fit to all data with RNAi and length of starvation as fixed effects and biological replicate as a random effect. ***interaction p-value < 0.001. (A-C and E-I) Cross bars reflect the mean (J) Model of intergenerational modification of IIS and its consequences on organismal physiology.

We used a *gld-1* mutant to further investigate the effect of IIS on germ cell tumor proliferation. 8 d L1 starvation increased the size of proximal tumors in this differentiation-defective mutant (Fig. S2C and D), as before (Fig. 2C and D). *daf-2* RNAi suppressed the effect of starvation, decreasing tumor size, in *daf-16*-dependent fashion (Fig. S2C and D). These results suggest that reduction of IIS after extended L1 starvation inhibits the hyperproliferative state of germ cells induced by starvation.

In addition to reduced brood size, extended L1 starvation results in production of abnormally shaped, small embryos with reduced hatching efficiency (*2*). Reduction of IIS by *daf-2* mutation or RNAi suppressed production of abnormal embryos, and it increased brood size following 8 d L1 starvation (Fig. 4H and I). These results further demonstrate that the starvation-induced germline abnormalities we have described compromise reproductive success, presumably reducing organismal fitness, and they show that reducing IIS following starvation increases reproductive success.

We have shown that early-life starvation leads to developmental abnormalities including tumors in the reproductive system of *C. elegans*, and that these abnormalities reduce reproductive success (Fig. 4J). Our results suggest that extended starvation followed by AL feeding causes hyperproliferation of germ cells, disrupting regulation of germ cell development and leading to tumor formation (*5, 6*). Remarkably, maternal DR mitigates these pathological consequences of starvation. We show that maternal DR reduces progeny IIS, and that somatic reduction of IIS following starvation suppresses germ cell hyperproliferation and development of abnormalities via activation of *daf-16*/FoxO and *skn-1*/Nrf, increasing reproductive success. These results add to an expanding body of work in *C. elegans* demonstrating intergenerational effects of the environment mediated by IIS (*4, 28-30*). Insulin signaling is frequently altered in mammalian models of maternal dietary effects on offspring (*31*), suggesting conservation of a central role of IIS in mediating physiological trade-offs between generations.

## Acknowledgments

We thank the Duke University School of Medicine and the Center for Genomic and Computational Biology for use of the Sequencing and Genomic Technologies core resource, which provided RNA sequencing service. Yossi Capua, Amanda Fry, and Theadora Tolkin for generating and validating *naSi2*. Some strains were provided by the CGC, which is funded by NIH Office of Research Infrastructure Programs (P40 OD010440). This work was funded by the National Institutes of Health (LRB, R01GM117408; EJAH, R01GM061706). JMJ, JDH, and LRB conceived and interpreted the experiments. JMJ and LRB wrote the manuscript. JMJ, JDH, REWK, AL, RG, and EAB conducted the experiments. JMJ, AKW, and CSM analyzed RNAseq data. EJAH provided an invaluable reagent and germline biology expertise. Authors declare no competing interests. RNAseq data set can be found at GEO using accession number GSE114271. Figure 1A adapted from *Caenorhabditis elegans hermaphrodite adult-en.svg* from Wikimedia Commons by K. D. Schroeder (License: CC-BY-SA 3.0).

## Supplementary Materials

## Materials and Methods

### Worm maintenance

Worms were maintained under standard laboratory conditions at 20°C unless otherwise noted. Animals were faithfully passaged without dietary restriction (thinning of the *E. coli* lawn) or starvation for many generations (greater than five) prior to commencing experiments unless otherwise noted.

### Strains used in this study

N2 (Bristol), GC1171 naSi2[pGC550(P*mex-5*<mCherry::H2B::*nos-2* 3’UTR<GFP::H2B::*nos-2* 3’UTR - *unc-119*(+))] *unc-119*(ed3), DP132 edIs6, OP354 *unc-119*(tm4063) III; wgIs354, CB1370 *daf-2(e1370)*, SPC167 dvIs19; *skn-1(lax120)*, SPC168 dvIs19; *skn-1(lax188)*, CF1038 *daf-16(mu86)*, NK1228 qyIs288; *daf-16(mu86)*; *unc-119(ed4)*, NK1229 qyIs290; *daf-16(mu86)*; *unc-119(ed4)*, NK1231 qyIs291; *daf-16(mu86)*; *unc-119(ed4)*, NK1233 qyIs293; *daf-16(mu86)*; *unc-119(ed4)*, NL2550 *ppw-1(pk2505)*, NL2098 *rrf-1(pk1417)*, JK3025 *gld-1(q485)*/hT2 [*bli-4(e937) let-?(*q782) qIs48], GC833 *glp-1(ar202)*, JK1505 *unc-32(e189) glp-1(e2072)*/eT1, and wild-type isolates: CB4856, ED3077, and JU561.

### mex-5 reporter construction

The *naSi2* allele results is an insertion of sequences from the plasmid pGC550 by MosSCI into strain EG4322(*32*). The insertion contains C. briggsae *unc-119* sequences and an FRT-flanked *mex-5*p::mCherry::H2B:*nos-2 3’* cassette followed by H2B::GFP. *naSi2* expresses mCherry::H2B in all germ cells. pGC550 was generated by a multisite LR reaction performed between pJA252 (gift from Julie Ahringer, Addgene plasmid # 21512, (*33*)), pGC544 [containing an FRT site (“<”) <*mCherry::H2B:: nos-2 3’UTR* sequences and inserted into pDONR-2221 by BP reaction], pGC545 [containing *Pmex-5<mCherry:: nos-2 3’UTR<GFP::nos-2 3’UTR*], and pCFJ150 - pDESTttTi5605[R4-R3] (gift from Erik Jorgensen, Addgene plasmid # 19329, (*32*)). All plasmids and sequences are available from Addgene or upon request.

### Recovery from Starvation Assay

*C. elegans’* embryos were prepared with sodium hypochlorite treatment, washed several times, and allowed to hatch in S-basal without additions. Animals were arrested for one or eight days (unless otherwise noted) prior to being plated on lawns of OP50 or HT115 *E. coli*. Animals were incubated at 20°C unless otherwise noted and then assayed after three or four days of development.

### Germline Microscopy and Scoring of Germline Abnormalities

Following recovery, adults were picked at random into 25 mM sodium azide on 2% noble agar pads. The worms were then examined using Nomarski microscopy on an AxioImager compound microscope (Zeiss) at 200-400x magnification. Animals with atypical gonads were scored as abnormal. A majority of the animals scored as abnormal had variations of proximal tumors that varied in the extent to which the germ cells were differentiated ranging from completely undifferentiated and *mex-5*^+^ to highly differentiated and *mex-5^-^*. However, starved animals also developed a variety of other abnormalities at varying rates including multiple, protruded, and extruded vulvas; gonad migration defects; egg-laying defects; endomitotic oocytes; and oocyte formation failures similar to those seen in *gld-2* and *gld-3* mutants (data not shown). When scoring *glp-1^gf^(ar202)*, dumpy individuals were not scored. Animals subjected to treatments and mutations which delayed development were scored later (up to 5 d after initiating recovery) to maximize chances of visualizing abnormal germline development. Animals were censored if they were delayed to the extent that the normality of their germline could not be determined at time of scoring. Homozygous *gld-1* mutant tumors were classified based on germ cell morphology for Figure 2D. For Figure 1E, each uterine mass was assessed for the presence or absence of P*elt-1* or P*unc-119*::GFP at 400-1000x.

### Gonad imaging

Worms were imaged at 400x magnification, using an AxioCam camera equipped with Zen software, on an AxioImager compound microscope (Zeiss). Images were merged using Fiji. Images were cropped and features were added using Inkscape.

### Determination of brood size

Animals were prepared as described above and plated onto the indicated bacterial lawn. After 48 hours of recovery, animals were singled onto plates seeded with the corresponding bacteria. Worms were transferred daily onto fresh plates until egg laying ceased and progeny were scored. Animals that could not be found were censored. For Fig. 1F, worms were inspected for germline abnormalities after 72 hours of recovery via dissection scope and otherwise scored as described.

### Abnormalities in progeny of AL and DR parents

Progeny from AL and DR parents were generated as previously described for use in this study (*4*). Briefly, following synchronization by brief L1 arrest, worms were cultured in S-complete at a density of 10 per mL with 25 mg/ml (AL) or 3.13 mg/ml (DR) *E. coli* HB101 at 20°C and 180 rpm for 96 hr. Progeny embryos were prepared by hypochlorite treatment.

### RNAseq in progeny from AL and DR parents

Progeny from AL and DR parents were cultured in S-basal for 24 hours so they hatch and enter L1 arrest, and then they were flash frozen in liquid nitrogen. RNA was prepared with Trizol (Invitrogen) according to the manufacturer’s instructions except that sand was included in homogenization. Libraries were prepared from 500 ng of total RNA and 12 PCR cycles using NEB Next Ultra RNA Library Preparation Kit (New England Biolabs). Sequencing was performed on an Illumina HiSeq 4000. Bowtie was used to map reads to the WS210 genome, including transcripts annotated in WS220 mapped back to WS210; (*34, 35*). HTSeq was used to generate count tables for each library (*36*). edgeR was used on count tables to determine differentially expressed genes. Detected genes were considered those with counts per million (CPM) > 1 across 3 libraries. This resulted in 13,439 genes for differential expression analysis. The “calcNormFactors” function was used and tagwise dispersion estimate was used for differential expression. Genes with a false-discovery rate of less than 0.1 were considered differentially expressed.

### daf-16-target gene signature analysis

A consensus background gene set was produced by overlapping all detected genes in the progeny of DR parents dataset with those reported by Tepper et al (*16*) using the R package gplots. Next, differentially expressed genes up- or down-regulated in the progeny of DR parents compared to AL parents (FDR < 0.1) were overlapped with high confidence (FDR < 0.001) *daf-16-*target genes as determined by Tepper et al (*16*). Hypergeometric p-values were determined using the Graeber lab calculator (http://systems.crump.ucla.edu/hypergeometric/index.php).

### Gene ontology analysis

Genes differentially expressed in the progeny of DR parents (114) were analyzed for enrichment of gene ontology terms against the 13439 detected genes (criteria detailed above) using GOrilla (*37*).

### GFP::DAF-16 localization in the progeny of AL and DR parents

Progeny of AL and DR parents were prepared as previously described for an N-terminal DAF-16 translational fusion reporter strain (NK1228). The extent of intestinal GFP localization was quantified at 400x or 1000x depending on stage using an AxioImager compound microscope (Zeiss).

### Starvation and DR recovery

Animals were starved as described above, except instead of recovery on plates animals were recovered in DR liquid culture as previously described prior to gonad assessment at 96 hr.

### Starvation to Dauer to AL recovery

Wild-type animals were starved and then recovered in dauer-forming conditions (*18*) (5 worms per µl and 1 mg/ml HB101 in S-complete at 20°C and 180 rpm) for 5 d then recovered on plates (AL). After animals reached early adulthood, their gonads were assessed for the presence or absence of germline abnormalities as described above.

### RNAi

Single colonies, grown on 100 µg/ml carbenicillin and 12.5 µg/ml tetracycline LB plates, were inoculated into LB starter cultures with the same antibiotics. These were used to inoculate larger cultures grown in 50 µg/ml carbenicillin. Cells from large cultures were spun down at 4°C and resuspended at a concentration of 25 mg/ml in 15% glycerol S-complete medium. Aliquots were frozen and thawed only once and seeded onto NGM plates containing 25 µg/ml carbenicillin and 1mM IPTG. For double RNAi experiments, equal volumes of RNAi suspension were mixed. Lawns were allowed to grow overnight at room temperature before adding worms. Empty vector (pAD12), *daf-2* (pAD48), and *daf-16* (pAD43) clones were gifts from Coleen Murphy. *skn-1* clones came from the Ahringer library.

### Data analysis

t-tests were performed using R or Microscoft Excel. Bartlett’s tests were performed to determine if variance could be pooled. Two-way ANOVA was performed using R. Plots were generated using the R package ggplot2 or Microsoft Excel. The R package plotrix was used to calculate SEM. The R package Linear and Nonlinear Mixed Effects Models “nmle” was used to compute the brood size interaction p-value (Fig. 4I).

---

**Fig. S1.**
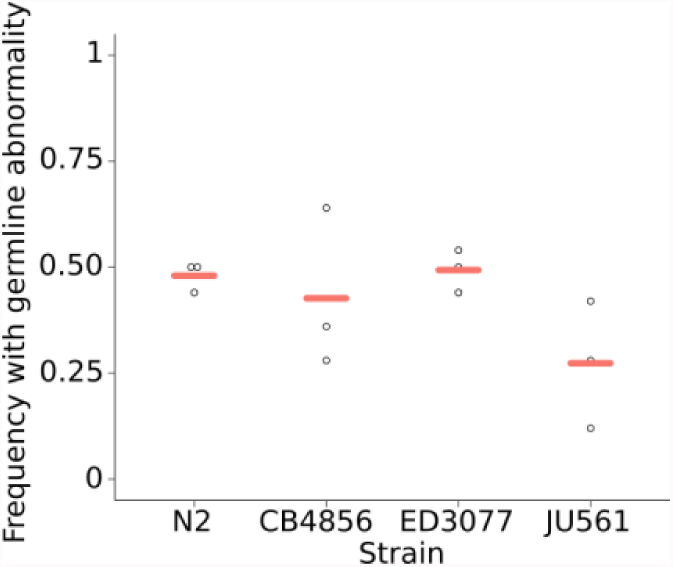
Abnormalities in various *C. elegans* wild-type isolates. Circles indicate biological replicates scoring at least 50 starved wild-type isolates per condition per replicate. Cross bars reflect the mean.

**Fig. S2.**
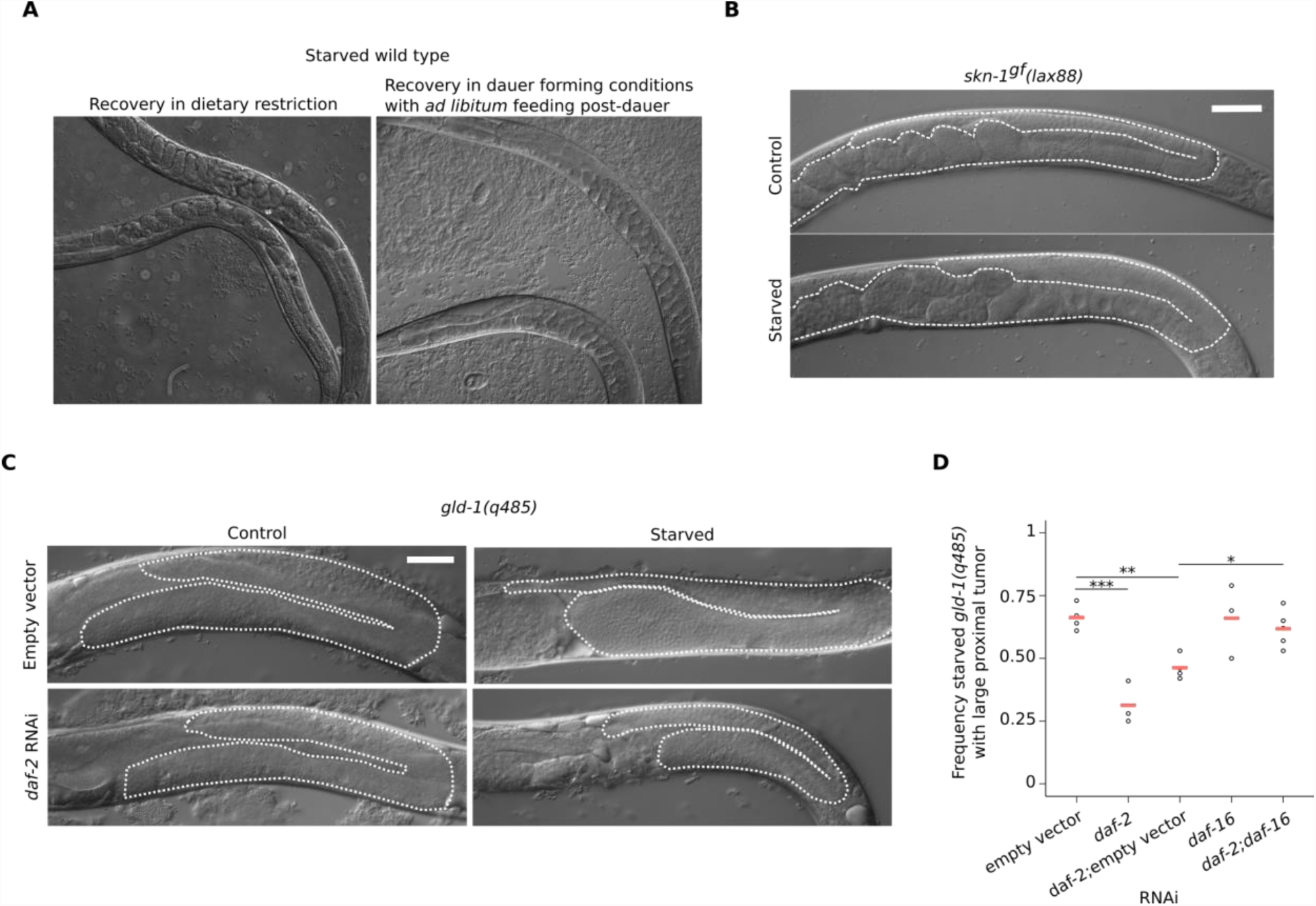
Reduced insulin/IGF signaling decreases germline proliferation following early-life starvation. (A-C) Representative image of early adult worms from indicated conditions. (B and C) Gonad and uterus are outlined with a dashed white line. Scale bar is 50 microns. (D) *daf-2* and *daf-16* RNAi in starved *gld-1(q485).* Circles indicate three biological replicates, scoring at least 30 worms per condition per replicate. Cross bars reflect the mean; *p < 0.05, ** < 0.01, *** < 0.001; t-test.

**Table S1.** Genes differentially expressed in the progeny of DR parents compared to AL parents during L1 arrest.

[See accompanying .csv file]

**Table S2.** GO terms from genes differentially expressed in the L1-arrested progeny of dietary restricted mothers.

| GO Term | Description | P-value | FDR q-value | Enrichment |
| --- | --- | --- | --- | --- |
| GO:0045087 | innate immune response | 1.7E-08 | 9.6E-05 | 8.2 |
| GO:0006955 | immune response | 2.0E-08 | 5.6E-05 | 8.1 |
| GO:0002376 | immune system process | 2.8E-08 | 5.4E-05 | 7.8 |
| GO:0006952 | defense response | 6.1E-08 | 8.7E-05 | 6.5 |
| GO:1905910 | negative regulation of dauer entry | 1.2E-05 | 1.3E-02 | 60.8 |
| GO:0006950 | response to stress | 5.1E-05 | 4.8E-02 | 3 |
| GO:1905909 | regulation of dauer entry | 5.3E-05 | 4.3E-02 | 38.7 |

**Table S3.**
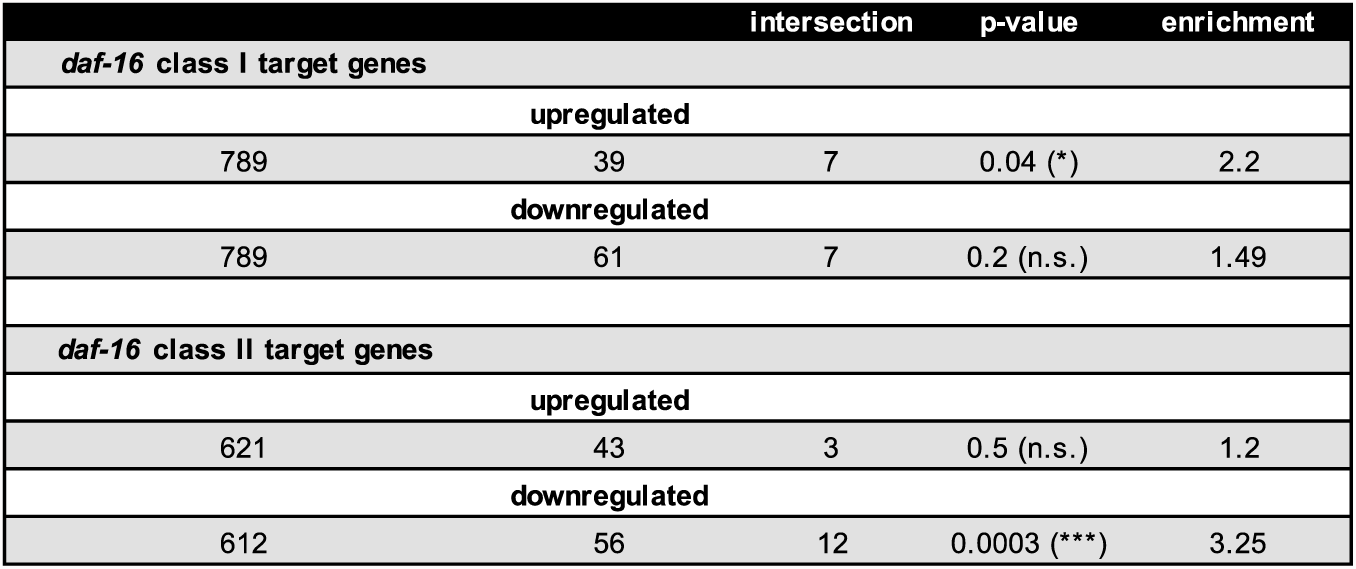
Intersection and hypergeometric p-values of *daf-16-*target genes and genes up or downregulated in the progeny of dietary restricted mothers.

|  |  | intersection | p-value | enrichment |
| --- | --- | --- | --- | --- |
| *daf-16* class I target genes |  |  |  |  |
| upregulated |  |  |  |  |
| 789 | 39 | 7 | 0.04 (*) | 2.2 |
| downregulated |  |  |  |  |
| 789 | 61 | 7 | 0.2 (n.s.) | 1.49 |
| *daf-16* class II target genes |  |  |  |  |
| upregulated |  |  |  |  |
| 621 | 43 | 3 | 0.5 (n.s.) | 1.2 |
| downregulated |  |  |  |  |
| 612 | 56 | 12 | 0.0003 (***) | 3.25 |

## References and Notes

1. L. R. Baugh, To grow or not to grow: nutritional control of development during Caenorhabditis elegans L1 arrest. Genetics 194, 539–555 (2013).

2. M. A. Jobson et al., Transgenerational Effects of Early Life Starvation on Growth, Reproduction, and Stress Resistance in Caenorhabditis elegans. Genetics 201, 201–212 (2015).

3. A. H. I. Lee, J. Kim, J. Yoshimoto, Y. You, Metabolic Rate Regulates L1 Longevity in C. elegans. PloS one 7, (2012).

4. J. D. Hibshman, A. Hung, L. R. Baugh, Maternal Diet and Insulin-Like Signaling Control Intergenerational Plasticity of Progeny Size and Starvation Resistance. PLoS Genet 12, e1006396 (2016).

5. M. McGovern, R. Voutev, J. Maciejowski, A. K. Corsi, E. J. Hubbard, A “latent niche” mechanism for tumor initiation. PNAS 106, 11617–11622 (2009).

6. R. Voutev, D. J. Killian, J. H. Ahn, E. J. Hubbard, Alterations in ribosome biogenesis cause specific defects in C. elegans hermaphrodite gonadogenesis. Dev Biol 298, 45–58 (2006).

7. J. Kimble, S. L. Crittenden, Germline proliferaton and its control. WormBook.org, (2005).

8. A. Pepper, D. J. Killian, E. J. Hubbard, Genetic Analysis of Caenorhabditis elegans glp-1 Mutants Suggests Receptor Interaction or Competition. Genetics 163, 115–132 (2003).

9. L. W. Berry, Westlund, B., Schedl, T., Germ-line tumor formation caused by activation of glp-1, a Caenorhabditis elegans member of the Notch family of receptors. Development 124, 925–936 (1997).

10. V. Kodoyianni, E. M. Maine, J. Kimble, Molecular Basis of Loss-of-Function Mutations in the glp-1 Gene of Caenorhabitis elegans. Molecular Biology of the Cell 3, 1199–1213 (1992).

11. R. Francis, M. K. Barton, J. Kimble, T. Schedl, gld-1, a Tumor Suppressor Gene Required for Oocyte Development in Caenorhabditis elegans. Genetics, 579–606 (1995).

12. J. M. Pinkston, D. Garigan, M. Hansen, C. Kenyon, Mutations That Increase the Life Span of C. elegans Inhibit Tumor Growth. Science 313, (2006).

13. T. L. Gumienny, E. Lambie, E. Hartwieg, H. R. Horvitz, M. O. Hengartner, Genetic control of programmed cell death in the Caenorhabditis elegans hermaphrodite germline. Development 126, 1011–1022 (1999).

14. W. Mair, S. H. Panowski, R. J. Shaw, A. Dillin, Optimizing dietary restriction for genetic epistasis analysis and gene discovery in C. elegans. PLoS One 4, e4535 (2009).

15. C. T. Murphy, P. J. Hu, Insulin/insulin-like growth factor signaling in C. elegans. WormBook, 1–43 (2013).

16. R. G. Tepper, Ashraf, J., Kaletsky, R., Kleemann, G., Murphy, C. T., Bussemaker, H. J., PQM-1 complements DAF-16 as a key transcriptional regulator of DAF-2-mediated development and longevity. Cell 154, 676–690 (2013).

17. Henderson S. T., T. E. Johnson, daf-16 integrates developmental and environmental inputs to mediate aging in the nematode Caenorhabditis elegans. Current Biology 11, 1975–1980 (2001).

18. L. R. Baugh, N. Kurhanewicz, P. W. Sternberg, Sensitive and precise quantification of insulin-like mRNA expression in Caenorhabditis elegans. PLoS One 6, e18086 (2011).

19. P. J. Hu, Dauer. WormBook, 1–19 (2007).

20. C. Kumsta, M. Hansen, C. elegans rrf-1 mutations maintain RNAi efficiency in the soma in addition to the germline. PLoS One 7, e35428 (2012).

21. O. Tijsterman M., K.L., Thijssen, K., Plasterk, R.H.A., PPW-1, a PAZ/PIWI Protein Required for Efficient Germline RNAi, Is Defective in a Natural Isolate of C. elegans. Current Biology 12, 1535–1540 (2002).

22. R. E. Kaplan, L. R. Baugh, L1 arrest, daf-16/FoxO and nonautonomous control of post-embryonic development. Worm 5, e1175196 (2016).

23. R. E. Kaplan, Chen, Y., Moore, B. T., Jordan, J. M., Maxwell, C. S., Schindler, A. J., Baugh, L. R., dbl-1/TGF-beta and daf-12/NHR Signaling Mediate Cell-Nonautonomous Effects of daf-16/FOXO on Starvation-Induced Developmental Arrest. PLoS Genet 11, e1005731 (2015).

24. N. Libina, J. R. Berman, C. Kenyon, Tissue-Specific Activities of C. elegans DAF-16 in the Regulation of Lifespan. Cell 115, 489–502 (2003).

25. P. Zhang, M. Judy, S. J. Lee, C. Kenyon, Direct and indirect gene regulation by a life-extending FOXO protein in C. elegans: roles for GATA factors and lipid gene regulators. Cell Metab 17, 85–100 (2013).

26. D. Michaelson, D. Z. Korta, Y. Capua, E. J. Hubbard, Insulin signaling promotes germline proliferation in C. elegans. Development 137, 671–680 (2010).

27. Z. Qin, E. J. Hubbard, Non-autonomous DAF-16/FOXO activity antagonizes age-related loss of C. elegans germline stem/progenitor cells. Nat Commun 6, 7107 (2015).

28. N. O. Burton, Furuta, T., Webster, A. K., Kaplan, R. E., Baugh, L. R., Arur, S., Horvitz, H. R., Insulin-like signalling to the maternal germline controls progeny response to osmotic stress. Nat Cell Biol 19, 252–257 (2017).

29. A. Tauffenberger, J. A. Parker, Heritable transmission of stress resistance by high dietary glucose in Caenorhabditis elegans. PLoS Genet 10, e1004346 (2014).

30. S. Kishimoto, M. Uno, E. Okabe, M. Nono, E. Nishida, Environmental stresses induce transgenerationally inheritable survival advantages via germline-to-soma communication in Caenorhabditis elegans. Nat Commun 8, 14031 (2017).

31. O. J. Rando, R. A. Simmons, I’m eating for two: parental dietary effects on offspring metabolism. Cell 161, 93–105 (2015).

## References and notes

32. C. Frokjaer-Jensen et al., Single-copy insertion of transgenes in Caenorhabditis elegans. Nature Genetics 40, 1375–1383 (2008).

33. E. Zeiser, C. Frokjaer-Jensen, E. Jorgensen, J. Ahringer, MosSCI and gateway compatible plasmid toolkit for constitutive and inducible expression of transgenes in the C. elegans germline. PLoS One 6, e20082 (2011).

34. B. Langmead, C. Trapnell, M. Pop, S. L. Salzberg, Ultrafast and memory-efficient alignment of short DNA sequences to the human genome. Genome Biol 10, R25 (2009).

35. C. S. Maxwell, I. Antoshechkin, N. Kurhanewicz, J. A. Belsky, L. R. Baugh, Nutritional control of mRNA isoform expression during developmental arrest and recovery in C. elegans. Genome Res 22, 1920–1929 (2012).

36. S. Anders, P. T. Pyl, W. Huber, HTSeq--a Python framework to work with high-throughput sequencing data. Bioinformatics 31, 166–169 (2015).

37. E. Eden, R. Navon, I. Steinfeld, D. Lipson, Z. Yakhini, GOrilla: a tool for discovery and visualization of enriched GO terms in ranked gene lists. BMC Bioinformatics 10, 48 (2009).

